## Supplementary Information for "Escape mutations circumvent a tradeoff between resistance to a beta-lactam and a beta-lactamase inhibitor"

[1] Faculty of Biology, Technion-Israel Institute of Technology, Haifa, Israel. [2] Lorry I. Lokey Interdisciplinary Center for Life Sciences and Engineering, Technion-Israel Institute of Technology, Haifa, Israel. [3] Roche Pharma Research and Early Development, Immunology, Infectious Diseases, and Ophthalmology, Roche Innovation Center Basel, F. Hoffmann-La Roche Ltd, Grenzacherstrasse 124, 4070 Basel, Switzerland.

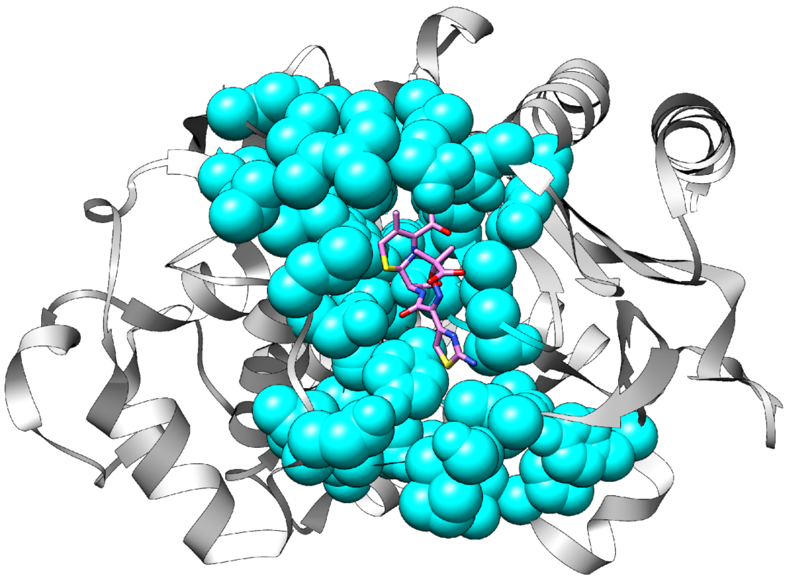

**Supplementary Figure 1. AmpC pocket.** Protein is shown in grey ribbon, residues in the pocket of AmpC are shown in cyan spheres, while the beta-lactam antibiotic ceftazidime is shown in sticks and colored by atom (C atoms colored pink, N blue, O red and S yellow).

**a**

**b**

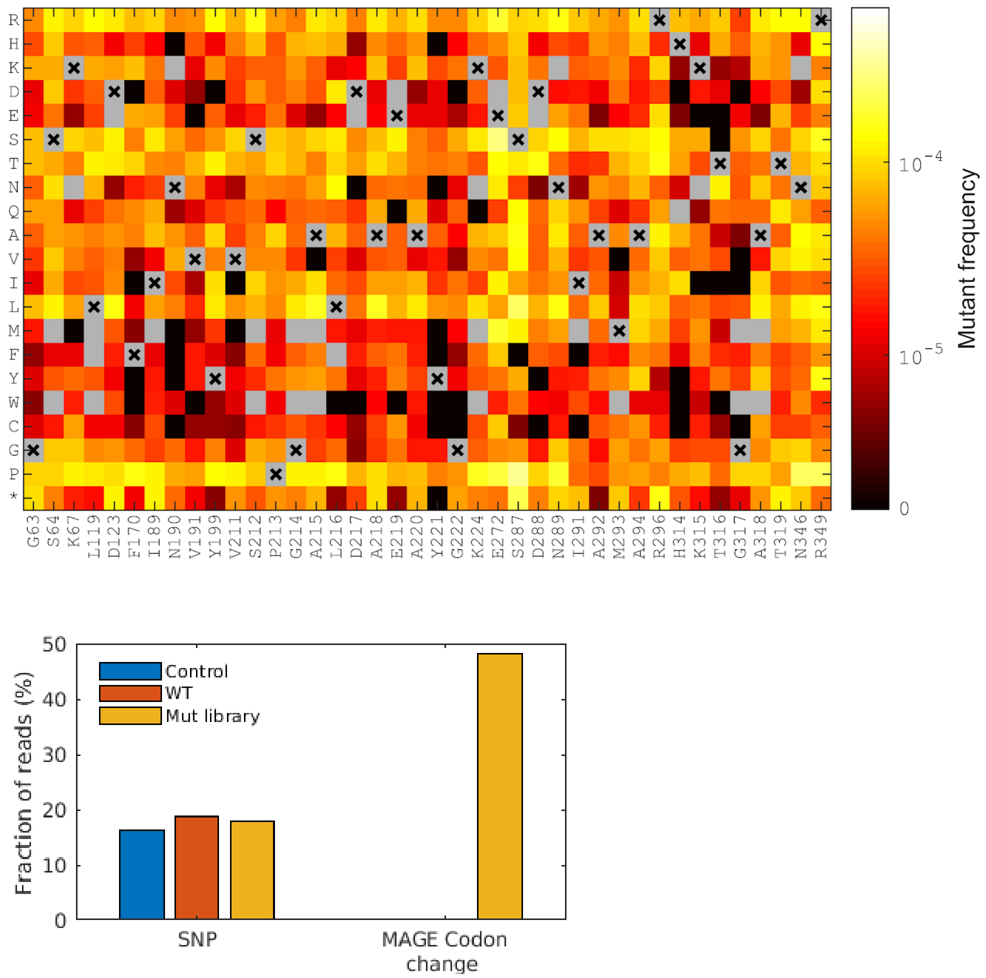


**Supplementary Figure 2. Mutants abundance in the mutants library. a.** Mutants frequency in the mutant library as identified by high-throughput sequencing of regions in the *ampC* gene. Since the mutated codons in the library are at least two nucleotides away from the original codon, not all amino acid substitutions are accessible. Non-accessible substitutions are marked in gray and the original amino acid with an X. The frequency data presents 40 of the 44 mutated residues excluding positions 148, 150, 151, and 152 which lie in the middle of the amplicon and were not well sequenced. Overall, the mean frequency of mutants is $5.3 \times 10^{-3}$ % ± $4.2 \times 10^{-3}$ % of the reads. **b.** Reads that contain one SNP (a single nucleotide change compared to the wild type sequence) as well as reads that contain mutations that resembles the designed MAGE mutations were identified from high-throughput sequencing of bases 768-999 in the *ampC* gene. The two types of mutations were identified in WT *ampC* strain and in the MAGE mutant library. Both strains were selected on a high drug concentration. In addition, mutations were identified in a control strain that was not selected on antibiotics (Methods). While all strains show similar frequency of SNPs, MAGE mutants appear mainly in the mutant library (48%), while very rare in the WT libraries (0.3%) and not at all in the control library.

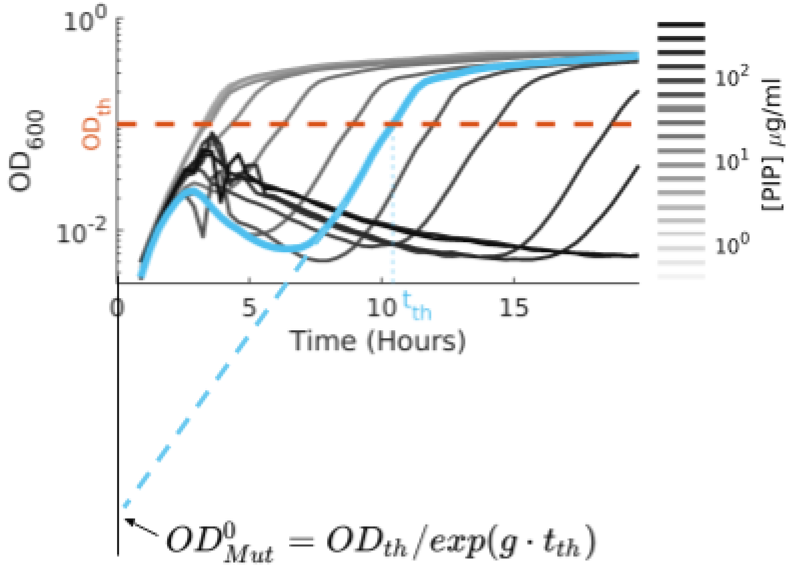


**Supplementary Figure 3. Calculating the initial number of viable mutants from growth measurements of the mutant library.** Growth of the mutant library exposed to increased concentration of PIP (black to light gray lines). To calculate the initial density of viable bacteria $OD^0_{Res}$ we extrapolate the exponential growth phase of the culture to calculate its OD at time 0 (dashed cyan line). Since bacterial density during exponential growth in rate g is $OD_{(t)} = OD_0 \cdot exp(g \cdot t)$ the initial density of viable cells is $OD^0_{Res} = OD_{th}/exp(g \cdot t_{th})$

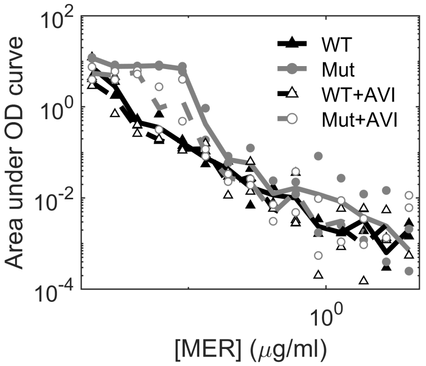


**Supplementary Figure 4. *ampC* overexpression does not increase bacterial resistance to meropenem.** Bacterial growth of the mutant library and the WT strain, as estimated by the area under the OD curve,, under stress of increasing MER concentrations with and without AVI. Growth of all strains was similar, implying that AmpC does not confer resistance to MER even when mutated.

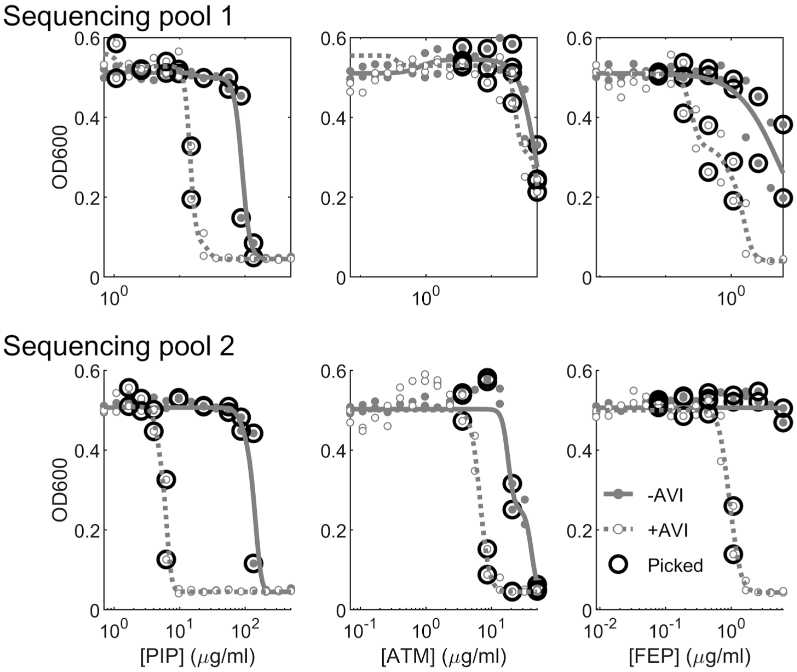


**Supplementary Figure 5. A subset of mutant libraries cultures selected on gradients of antibiotics with and without avibactam was picked for high-throughput sequencing.** Mutant libraries comprised of mutants with substitutions in the region of sequencing pool 1 (top, nucleotides 141-624) and sequencing pool 2 (bottom, nucleotides 768-999) were cultured in LB on gradients of three different drugs with and without avibactam. A subset of the cultures (circles) was picked for genetic analysis.

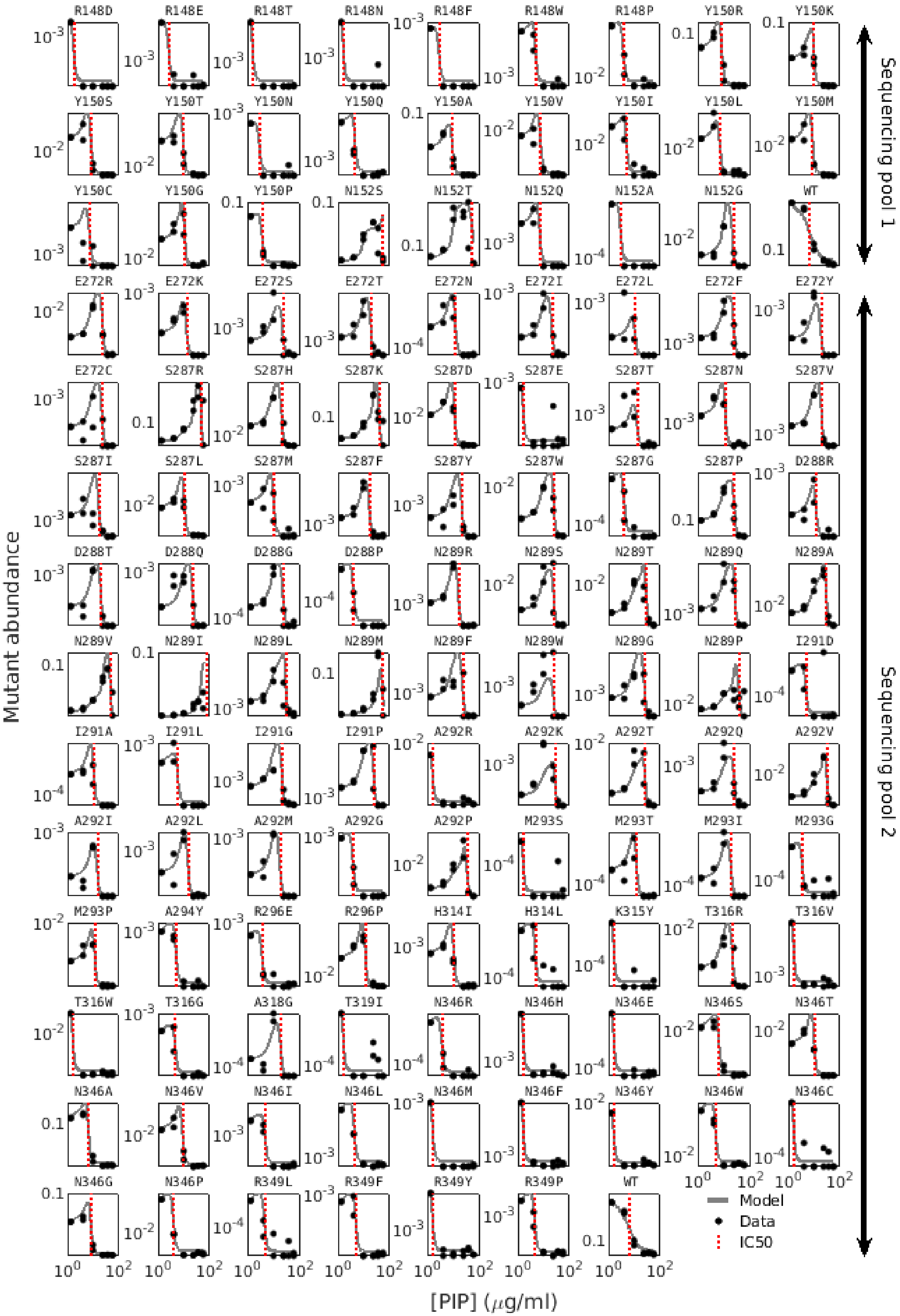

R148D R148E R148T R148N R148F R148W R148P Y150R Y150K
Y150S Y150T Y150N Y150Q Y150A Y150V Y150I Y150L Y150M
Y150C Y150G Y150P N152S N152T N152Q N152A N152G WT
E272R E272K E272S E272T E272N E272I E272L E272F E272Y
E272C S287R S287H S287K S287D S287E S287T S287N S287V
S287I S287L S287M S287F S287Y S287W S287G S287P D288R
D288T D288Q D288G D288P N289R N289S N289T N289Q N289A
N289V N289I N289L N289M N289F N289W N289G N289P I291D
I291A I291L I291G I291P A292R A292K A292T A292Q A292V
A292I A292L A292M A292G A292P M293S M293T M293I M293G
M293P A294Y R296E R296P H314I H314L K315Y T316R T316V
T316W T316G A318G T319I N346R N346H N346E N346S N346T
N346A N346V N346I N346L N346M N346F N346Y N346W N346C
N346G N346P R349L R349F R349Y R349P WT
Mutant abundance
[PIP] (μg/ml)
Model
Data
IC50
Sequencing pool 1
Sequencing pool 2

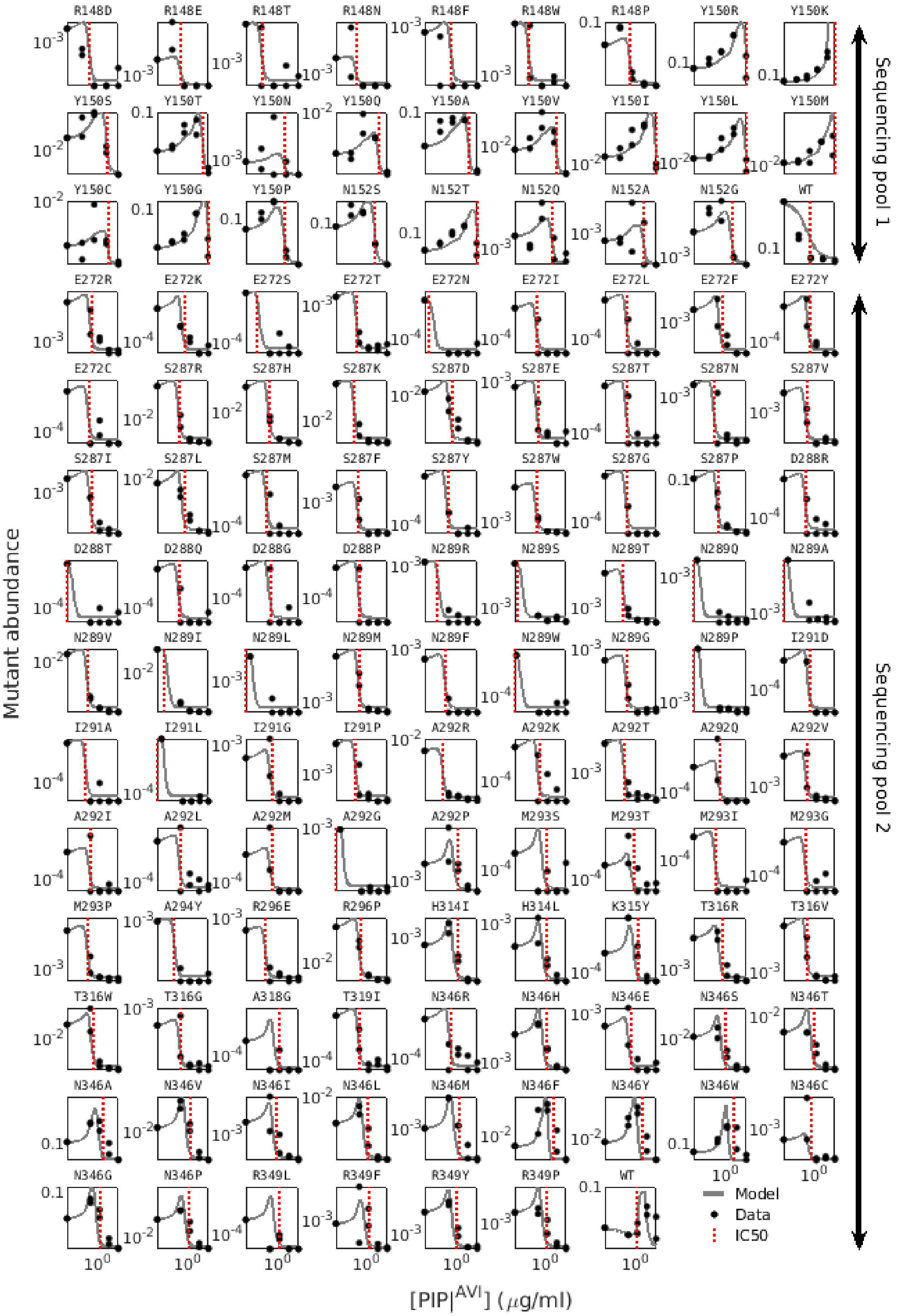
R148D
R148E
R148T
R148N
R148F
R148W
R148P
Y150R
Y150K
Y150S
Y150T
Y150N
Y150Q
Y150A
Y150V
Y150I
Y150L
Y150M
Y150C
Y150G
Y150P
N152S
N152T
N152Q
N152A
N152G
WT
E272R
E272K
E272S
E272T
E272N
E272I
E272L
E272F
E272Y
E272C
S287R
S287H
S287K
S287D
S287E
S287T
S287N
S287V
S287I
S287L
S287M
S287F
S287Y
S287W
S287G
S287P
D288R
D288T
D288Q
D288G
D288P
N289R
N289S
N289T
N289Q
N289A
N289V
N289I
N289L
N289M
N289F
N289W
N289G
N289P
I291D
I291A
I291L
I291G
I291P
A292R
A292K
A292T
A292Q
A292V
A292I
A292L
A292M
A292G
A292P
M293S
M293T
M293I
M293G
M293P
A294Y
R296E
R296P
H314I
H314L
K315Y
T316R
T316V
T316W
T316G
A318G
T319I
N346R
N346H
N346E
N346S
N346T
N346A
N346V
N346I
N346L
N346M
N346F
N346Y
N346W
N346C
N346G
N346P
R349L
R349F
R349Y
R349P
WT
Model
Data
IC50
Mutant abundance
[PIP|AVI] (μg/ml)
Sequencing pool 1
Sequencing pool 2

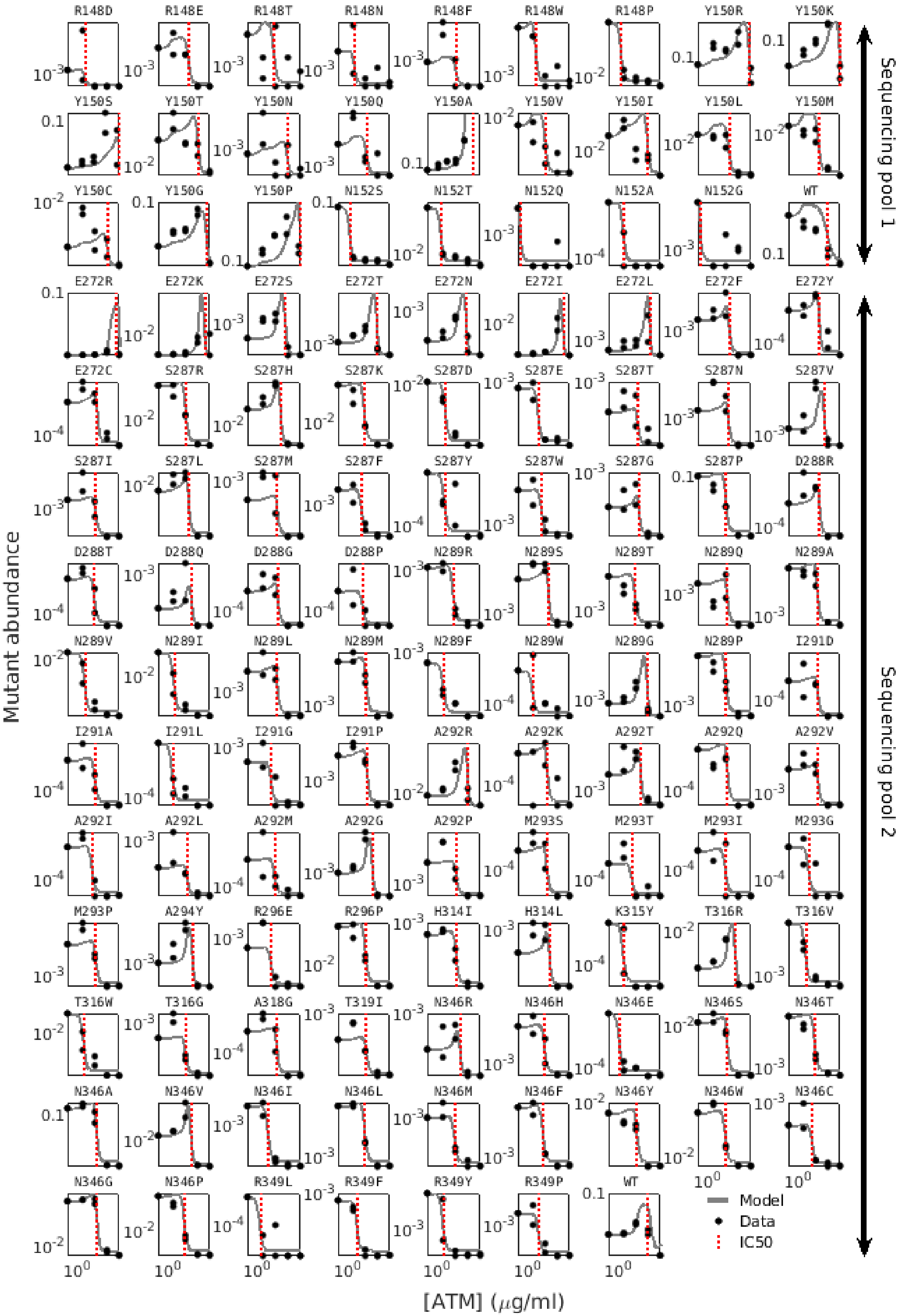
R148D
R148E
R148T
R148N
R148F
R148W
R148P
Y150R
Y150K
Y150S
Y150T
Y150N
Y150Q
Y150A
Y150V
Y150I
Y150L
Y150M
Y150C
Y150G
Y150P
N152S
N152T
N152Q
N152A
N152G
WT
E272R
E272K
E272S
E272T
E272N
E272I
E272L
E272F
E272Y
E272C
S287R
S287H
S287K
S287D
S287E
S287T
S287N
S287V
S287I
S287L
S287M
S287F
S287Y
S287W
S287G
S287P
D288R
D288T
D288Q
D288G
D288P
N289R
N289S
N289T
N289Q
N289A
N289V
N289I
N289L
N289M
N289F
N289W
N289G
N289P
I291D
I291A
I291L
I291G
I291P
A292R
A292K
A292T
A292Q
A292V
A292I
A292L
A292M
A292G
A292P
M293S
M293T
M293I
M293G
M293P
A294Y
R296E
R296P
H314I
H314L
K315Y
T316R
T316V
T316W
T316G
A318G
T319I
N346R
N346H
N346E
N346S
N346T
N346A
N346V
N346I
N346L
N346M
N346F
N346Y
N346W
N346C
N346G
N346P
R349L
R349F
R349Y
R349P
WT
Model
Data
IC50
Mutant abundance
[ATM] (μg/ml)
Sequencing pool 1
Sequencing pool 2

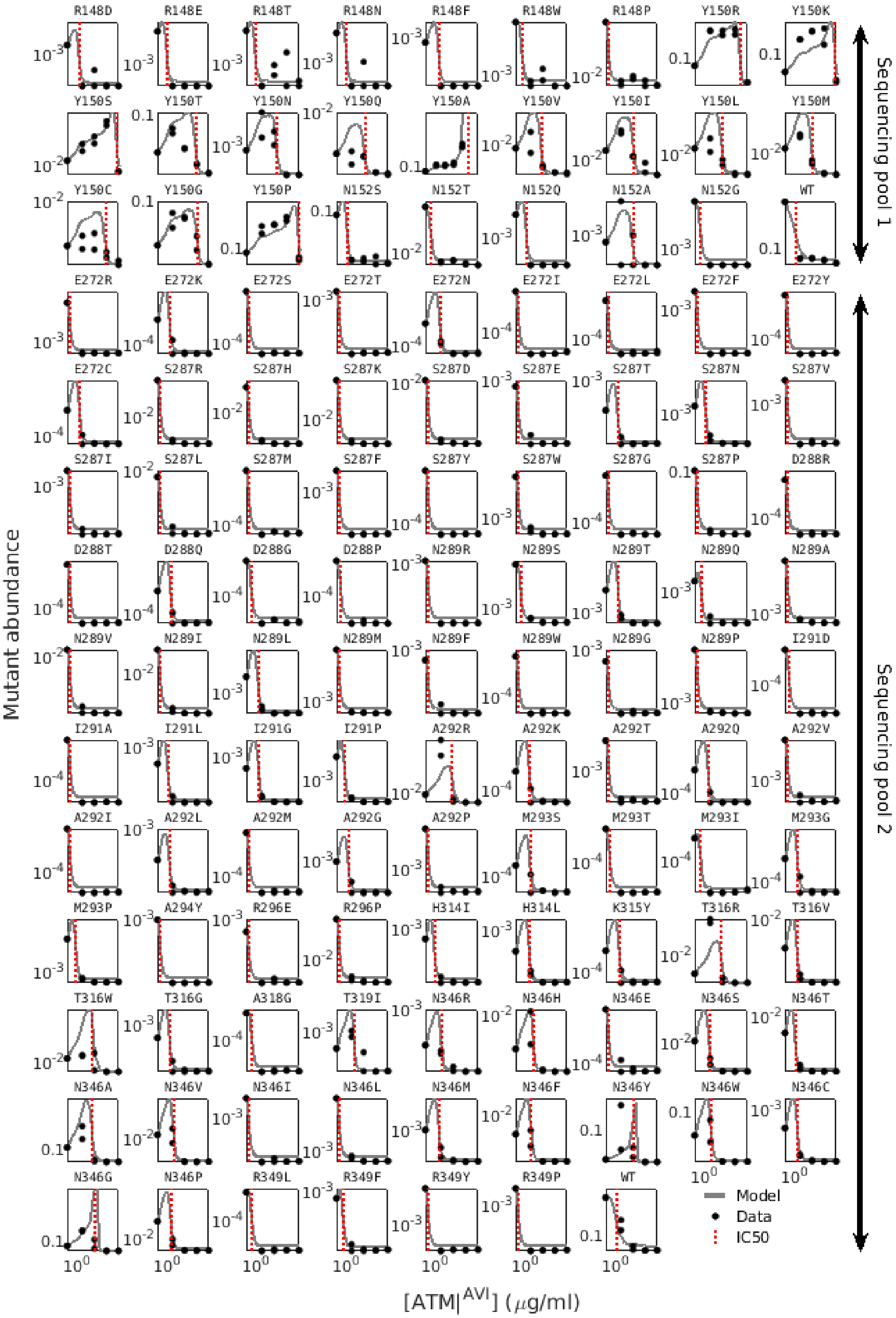
R148D R148E R148T R148N R148F R148W R148P Y150R Y150K
Y150S Y150T Y150N Y150Q Y150A Y150V Y150I Y150L Y150M
Y150C Y150G Y150P N152S N152T N152Q N152A N152G WT
E272R E272K E272S E272T E272N E272I E272L E272F E272Y
E272C S287R S287H S287K S287D S287E S287T S287N S287V
S287I S287L S287M S287F S287Y S287W S287G S287P D288R
D288T D288Q D288G D288P N289R N289S N289T N289Q N289A
N289V N289I N289L N289M N289F N289W N289G N289P I291D
I291A I291L I291G I291P A292R A292K A292T A292Q A292V
A292I A292L A292M A292G A292P M293S M293T M293I M293G
M293P A294Y R296E R296P H314I H314L K315Y T316R T316V
T316W T316G A318G T319I N346R N346H N346E N346S N346T
N346A N346V N346I N346L N346M N346F N346Y N346W N346C
N346G N346P R349L R349F R349Y R349P WT
Model
Data
IC50
Mutant abundance
[ATM|$^{AVI}$] ($\mu$g/ml)
Sequencing pool 1
Sequencing pool 2

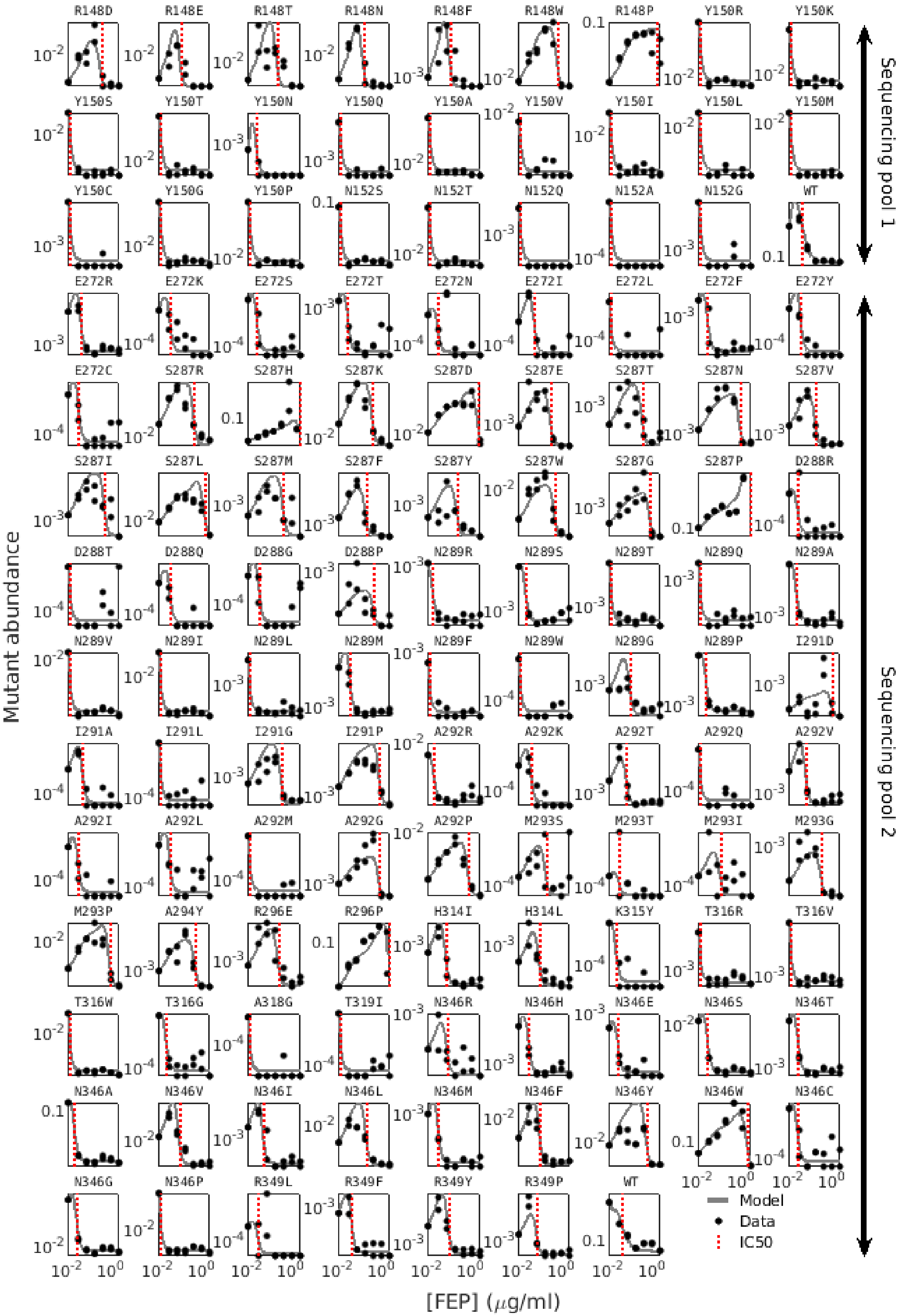

R148D R148E R148T R148N R148F R148W R148P Y150R Y150K
Y150S Y150T Y150N Y150Q Y150A Y150V Y150I Y150L Y150M
Y150C Y150G Y150P N152S N152T N152Q N152A N152G WT
E272R E272K E272S E272T E272N E272I E272L E272F E272Y
E272C S287R S287H S287K S287D S287E S287T S287N S287V
S287I S287L S287M S287F S287Y S287W S287G S287P D288R
D288T D288Q D288G D288P N289R N289S N289T N289Q N289A
N289V N289I N289L N289M N289F N289W N289G N289P I291D
I291A I291L I291G I291P A292R A292K A292T A292Q A292V
A292I A292L A292M A292G A292P M293S M293T M293I M293G
M293P A294Y R296E R296P H314I H314L K315Y T316R T316V
T316W T316G A318G T319I N346R N346H N346E N346S N346T
N346A N346V N346I N346L N346M N346F N346Y N346W N346C
N346G N346P R349L R349F R349Y R349P WT
Model
Data
IC50
Mutant abundance
[FEP] ($\mu$g/ml)
Sequencing pool 1
Sequencing pool 2

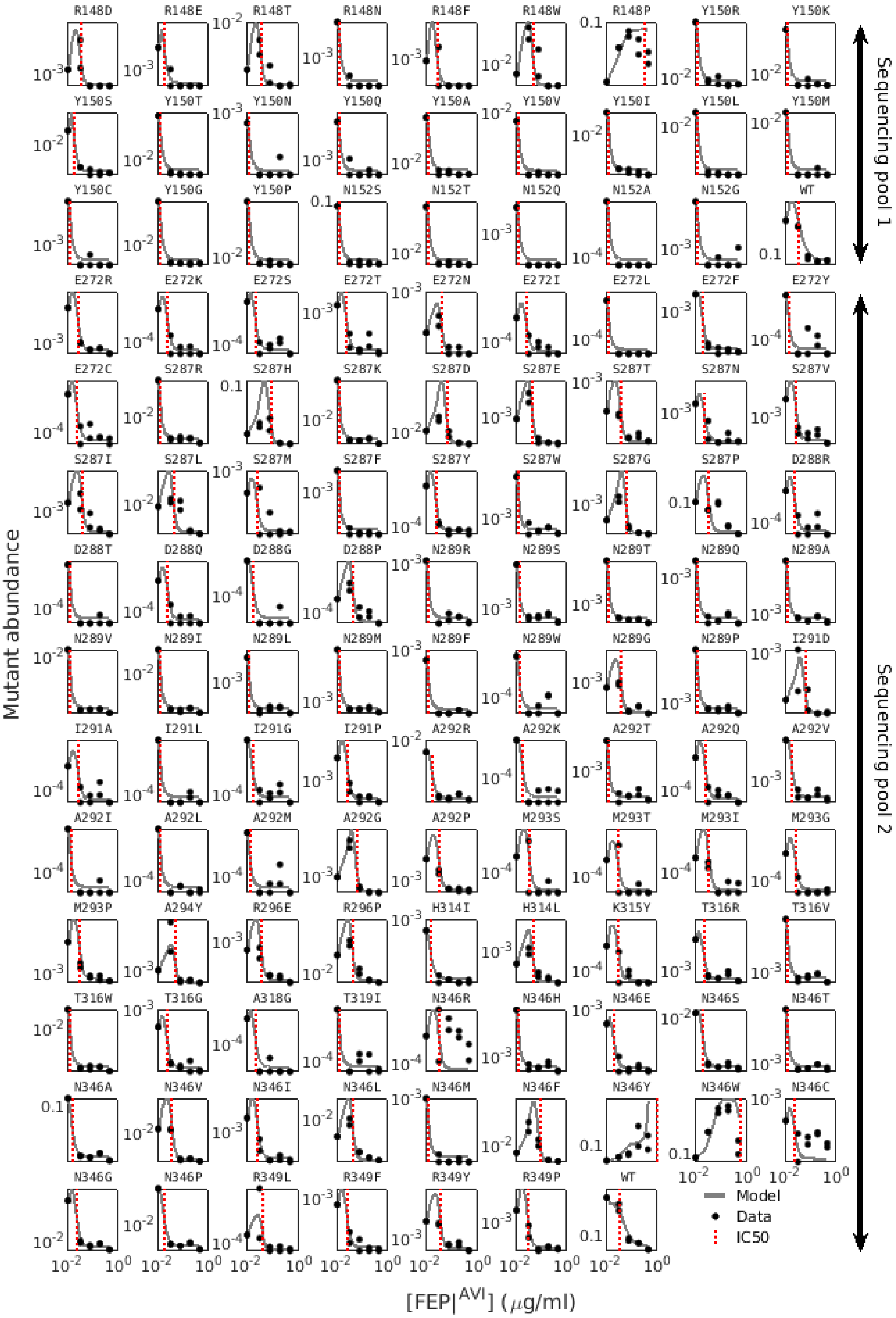

R148D
R148E
R148T
R148N
R148F
R148W
R148P
Y150R
Y150K
Y150S
Y150T
Y150N
Y150Q
Y150A
Y150V
Y150I
Y150L
Y150M
Y150C
Y150G
Y150P
N152S
N152T
N152Q
N152A
N152G
WT
E272R
E272K
E272S
E272T
E272N
E272I
E272L
E272F
E272Y
E272C
S287R
S287H
S287K
S287D
S287E
S287T
S287N
S287V
S287I
S287L
S287M
S287F
S287Y
S287W
S287G
S287P
D288R
D288T
D288Q
D288G
D288P
N289R
N289S
N289T
N289Q
N289A
N289V
N289I
N289L
N289M
N289F
N289W
N289G
N289P
I291D
I291A
I291L
I291G
I291P
A292R
A292K
A292T
A292Q
A292V
A292I
A292L
A292M
A292G
A292P
M293S
M293T
M293I
M293G
M293P
A294Y
R296E
R296P
H314I
H314L
K315Y
T316R
T316V
T316W
T316G
A318G
T319I
N346R
N346H
N346E
N346S
N346T
N346A
N346V
N346I
N346L
N346M
N346F
N346Y
N346W
N346C
N346G
N346P
R349L
R349F
R349Y
R349P
WT
Model
Data
IC50
Mutant abundance
[FEP|AVI] (μg/ml)
Sequencing pool 1
Sequencing pool 2

**Supplementary Figure 6. Mutants susceptibility to different drugs is estimated by fitting reads abundance to a mathematical model of inter-mutant competition.** The relative abundance ($RA_{Mut}$) of each mutant is estimated from high-throughput sequencing and used to calculate the 'mutant abundance', $RA_{Mut}\ x\ OD_{Culture}$ (Duplicate measurements, black dots). The abundance data was fitted to a dose response model to calculate the mutant resistance ($IC50$, red dashed line). based on inter-mutant competition (Methods, gray line).

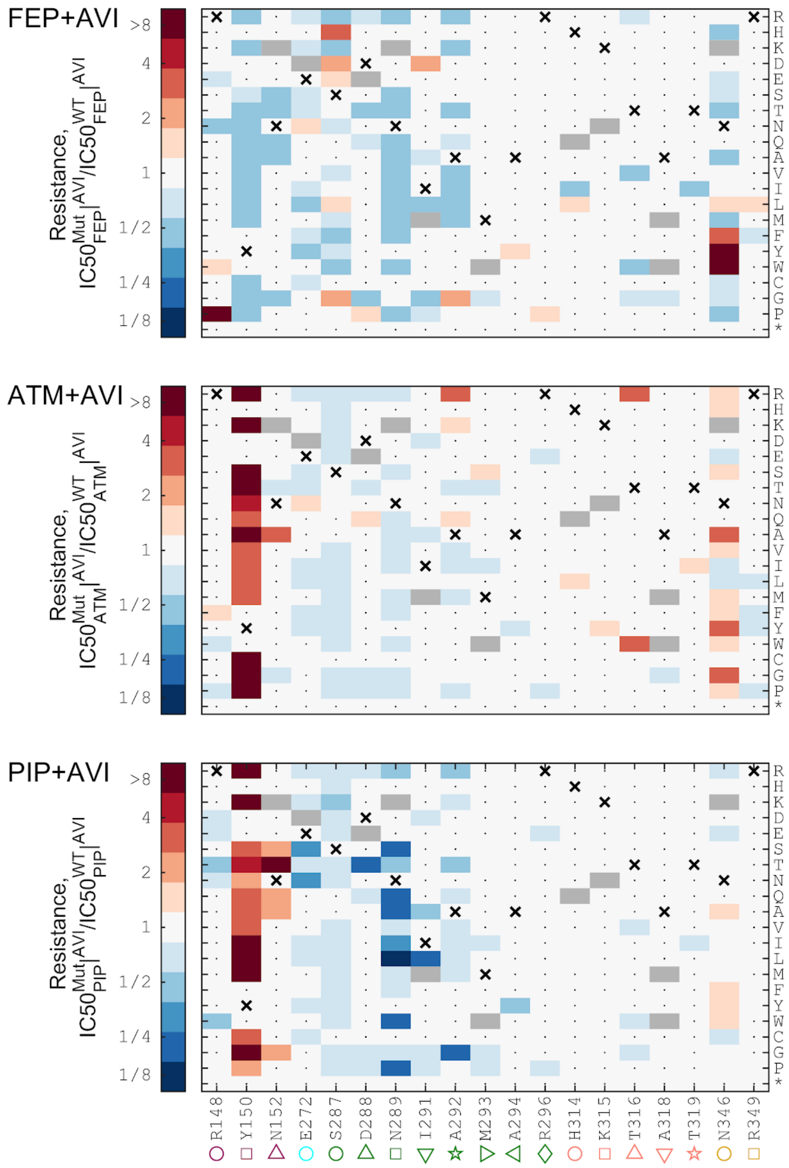


**Supplementary Figure 7. The resistance of all mutants to PIP, ATM, and FEP when co-applied with AVI as calculated from high-throughput sequencing data.** The relative resistance of all identified mutants to the beta-lactam drugs when supplemented with 0.25 $\mu$g/ml avibactam is calculated based on a mathematical model capturing inter-mutant competition (Methods).

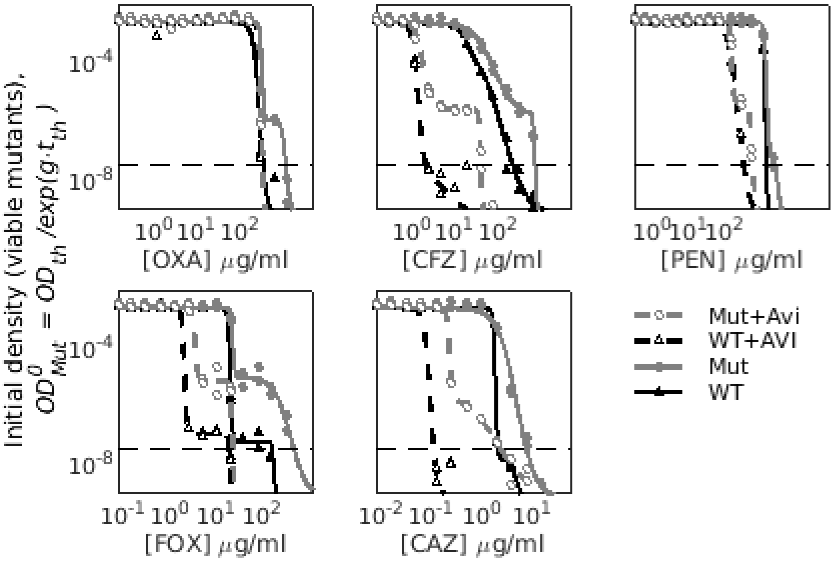


**Supplementary Figure 8. The initial density of viable mutants in the additional five drugs.** The initial density of viable mutants, $OD^0_{Res}$, is calculated from growth measurements for five beta-lactam drugs. While escape mutants exist for a narrow concentration range of cefazolin (CAZ), no escape mutations appear for the other drugs.

**Supplementary table 1. Oligos used in this study**

| | |
|---|---|
| ampC_sp1_int F 1 | TCGTCGGCAGCGTCAGATGTGTATAAGAGACAGCCCGTCACACAGCAAAC |
| ampC_sp1_int F 2 | TCGTCGGCAGCGTCAGATGTGTATAAGAGACAGCCGTCACACAGCAAACG |
| ampC_sp1_int F 3 | TCGTCGGCAGCGTCAGATGTGTATAAGAGACAGCGTCACACAGCAAACGTT |
| ampC_sp1_int R 1 | GTCTCGTGGGCTCGGAGATGTGTATAAGAGACAGAGCGGGCCATATCTTCA |
| ampC_sp1_int R 2 | GTCTCGTGGGCTCGGAGATGTGTATAAGAGACAGGCGGGCCATATCTTCAA |
| ampC_sp1_int R 3 | GTCTCGTGGGCTCGGAGATGTGTATAAGAGACAGCGGGCCATATCTTCAATG |
| ampC_sp2_int F 1 | TCGTCGGCAGCGTCAGATGTGTATAAGAGACAGGCAAAGCAATTTAAAACCCCT |
| ampC_sp2_int F 2 | TCGTCGGCAGCGTCAGATGTGTATAAGAGACAGAAAGCAATTTAAAACCCCTTG |
| ampC_sp2_int F 3 | TCGTCGGCAGCGTCAGATGTGTATAAGAGACAGAGCAATTTAAAACCCCTTGATAT |
| ampC_sp2_int R 1 | GTCTCGTGGGCTCGGAGATGTGTATAAGAGACAGCCGATGGAATTTTACTGTAGAGC |

| | |
|---|---|
| ampC_sp2_int R 2 | GTCTCGTGGGCTCGGAGATGTGTATAAGAGACAGCGATGGAATTTTACTGTAGAGCG |
| ampC_sp2_int R 3 | GTCTCGTGGGCTCGGAGATGTGTATAAGAGACAGGATGGAATTTTACTGTAGAGCGTT |
| Ampx_sp2_seq_gap F | GCGCTTCTATCAAAACTGGCAGCCTGCAT |
| Ampx_sp2_seq_gap R | GGTTGAGTTTGAGTGGCTGGAAGACACGA |
| nx_i5 | AATGATACGGCGACCACCGAGATCTACACXXXXXXXXTCGTCGGCAGCGTC |
| nx_i7 | CAAGCAGAAGACGGCATACGAGATXXXXXXXXGTCTCGTGGGCTCGG |
| MS_AmpC_G79_1_p1 | T*A*G* CGT CGC CAC CAA GCA CGC CAG TAA ATG TTT TGC TGA CCG ABN DTA ACT CAA ACA ACG TTT GCT GTG TGA CGG GCT GCT TTT TGG CGA |
| MS_AmpC_G79_2_p1 | T*A*G* CGT CGC CAC CAA GCA CGC CAG TAA ATG TTT TGC TGA CCG ABD NTA ACT CAA ACA ACG TTT GCT GTG TGA CGG GCT GCT TTT TGG CGA |
| MS_AmpC_S80_1_p1 | C*A*A* TAG CGT CGC CAC CAA GCA CGC CAG TAA ATG TTT TGC TGA CDN BAC CTA ACT CAA ACA ACG TTT GCT GTG TGA CGG GCT GCT TTT TGG |
| MS_AmpC_S80_2_p1 | C*A*A* TAG CGT CGC CAC CAA GCA CGC CAG TAA ATG TTT TGC TGA CDH NAC CTA ACT CAA ACA ACG TTT GCT GTG TGA CGG GCT GCT TTT TGG |
| MS_AmpC_K83_1_p1 | C*C*C* CTC GAG CAA TAG CGT CGC CAC CAA GCA CGC CAG TAA ATG TVN VGC TGA CCG AAC CTA ACT CAA ACA ACG TTT GCT GTG TGA CGG GCT |
| MS_AmpC_K83_2_p1 | C*C*C* CTC GAG CAA TAG CGT CGC CAC CAA GCA CGC CAG TAA ATG TVV NGC TGA CCG AAC CTA ACT CAA ACA ACG TTT GCT GTG TGA CGG GCT |
| MS_AmpC_L135_1_p2 | A*G*C* GCA GCA AGT CGC TTG AGG ATT TCA CCT CAT CCG GCA CCT GDN BTG GCA GGC CGC CAG CAG TGT AGG TTG CGA GAT GTA ATA GTG TGA |
| MS_AmpC_L135_2_p2 | A*G*C* GCA GCA AGT CGC TTG AGG ATT TCA CCT CAT CCG GCA CCT GDB NTG GCA GGC CGC CAG CAG TGT AGG TTG CGA GAT GTA ATA GTG TGA |
| MS_AmpC_D139_1_p2 | A*G*T* TTT GAT AGA AGC GCA GCA AGT CGC TTG AGG ATT TCA CCT CBN DCG GCA CCT GCA ATG GCA GGC CGC CAG CAG TGT AGG TTG CGA GAT |
| MS_AmpC_D139_2_p2 | A*G*T* TTT GAT AGA AGC GCA GCA AGT CGC TTG AGG ATT TCA CCT CBV NCG GCA CCT GCA ATG GCA GGC CGC CAG CAG TGT AGG TTG CGA GAT |
| MS_AmpC_R164_1_p3 | C*A*G* CCA GTG CGC CGA ACA AAC CGA TAC TGG AGT TGG CAT ACA GBN HTT GTG TTC CTG GAG CCC ATG CAG GCT GCC AGT TTT GAT AGA AGC |
| MS_AmpC_R164_2_p3 | C*A*G* CCA GTG CGC CGA ACA AAC CGA TAC TGG AGT TGG CAT ACA GBD NTT GTG TTC CTG GAG CCC ATG CAG GCT GCC AGT TTT GAT AGA AGC |

| | |
|---|---|
| MS_AmpC_L165_1_p3 | T*C*A* CAG CCA GTG CGC CGA ACA AAC CGA TAC TGG AGT TGG CAT ADN HAC GTT GTG TTC CTG GAG CCC ATG CAG GCT GCC AGT TTT GAT AGA |
| MS_AmpC_L165_2_p3 | T*C*A* CAG CCA GTG CGC CGA ACA AAC CGA TAC TGG AGT TGG CAT ADB NAC GTT GTG TTC CTG GAG CCC ATG CAG GCT GCC AGT TTT GAT AGA |
| MS_AmpC_Y166_1_p3 | G*C*T* TCA CAG CCA GTG CGC CGA ACA AAC CGA TAC TGG AGT TGG CBN BCA GAC GTT GTG TTC CTG GAG CCC ATG CAG GCT GCC AGT TTT GAT |
| MS_AmpC_Y166_2_p3 | G*C*T* TCA CAG CCA GTG CGC CGA ACA AAC CGA TAC TGG AGT TGG CBV NCA GAC GTT GTG TTC CTG GAG CCC ATG CAG GCT GCC AGT TTT GAT |
| MS_AmpC_N168_1_p3 | C*A*G* ACG GCT TCA CAG CCA GTG CGC CGA ACA AAC CGA TAC TGG AHN VGG CAT ACA GAC GTT GTG TTC CTG GAG CCC ATG CAG GCT GCC AGT |
| MS_AmpC_N168_2_p3 | C*A*G* ACG GCT TCA CAG CCA GTG CGC CGA ACA AAC CGA TAC TGG AHV NGG CAT ACA GAC GTT GTG TTC CTG GAG CCC ATG CAG GCT GCC AGT |
| MS_AmpC_F186_1_p4 | T*G*A* GTT TGA GTG GCT GGA AGA CAC GAG TTT GCA TCG CCT GCT CBN BAC TCA AAC CAG ACG GCT TCA CAG CCA GTG CGC CGA ACA AAC CGA |
| MS_AmpC_F186_2_p4 | T*G*A* GTT TGA GTG GCT GGA AGA CAC GAG TTT GCA TCG CCT GCT CBB NAC TCA AAC CAG ACG GCT TCA CAG CCA GTG CGC CGA ACA AAC CGA |
| MS_AmpC_I205_1_p5 | G*A*T* ATC CCC AGG CGT AAT TCT TTT CTT CTG CGG GCG GTA CAT TBN VCC ACG TAT GGT TGA GTT TGA GTG GCT GGA AGA CAC GAG TTT GCA |
| MS_AmpC_I205_2_p5 | G*A*T* ATC CCC AGG CGT AAT TCT TTT CTT CTG CGG GCG GTA CAT TBB NCC ACG TAT GGT TGA GTT TGA GTG GCT GGA AGA CAC GAG TTT GCA |
| MS_AmpC_N206_1_p5 | C*G*C* GAT ATC CCC AGG CGT AAT TCT TTT CTT CTG CGG GCG GTA CBN VAA TCC ACG TAT GGT TGA GTT TGA GTG GCT GGA AGA CAC GAG TTT |
| MS_AmpC_N206_2_p5 | C*G*C* GAT ATC CCC AGG CGT AAT TCT TTT CTT CTG CGG GCG GTA CBV NAA TCC ACG TAT GGT TGA GTT TGA GTG GCT GGA AGA CAC GAG TTT |
| MS_AmpC_V207_1_p5 | C*T*T* CGC GAT ATC CCC AGG CGT AAT TCT TTT CTT CTG CGG GCG GVN DAT TAA TCC ACG TAT GGT TGA GTT TGA GTG GCT GGA AGA CAC GAG |
| MS_AmpC_V207_2_p5 | C*T*T* CGC GAT ATC CCC AGG CGT AAT TCT TTT CTT CTG CGG GCG GVB NAT TAA TCC ACG TAT GGT TGA GTT TGA GTG GCT GGA AGA CAC GAG |
| MS_AmpC_Y215_1_p5 | C*A*G* GCG AAA CAT GCA CTG CCT TAC CTT CGC GAT ATC CCC AGG CHN BAT TCT TTT CTT CTG CGG GCG GTA CAT TAA TCC ACG TAT GGT TGA |
| MS_AmpC_Y215_2_p5 | C*A*G* GCG AAA CAT GCA CTG CCT TAC CTT CGC GAT ATC CCC AGG CHV NAT TCT TTT CTT CTG CGG GCG GTA CAT TAA TCC ACG TAT GGT TGA |

| | |
|---|---|
| MS_AmpC_V227_1_p6 | T*C*G* ACT TCA CAC CAT AAG CTT CAG CAT CTA ACG CCC CAG GCG ABN DAT GCA CTG CCT TAC CTT CGC GAT ATC CCC AGG CGT AAT TCT TTT |
| MS_AmpC_V227_2_p6 | T*C*G* ACT TCA CAC CAT AAG CTT CAG CAT CTA ACG CCC CAG GCG ABB NAT GCA CTG CCT TAC CTT CGC GAT ATC CCC AGG CGT AAT TCT TTT |
| MS_AmpC_S228_1_p6 | T*G*G* TCG ACT TCA CAC CAT AAG CTT CAG CAT CTA ACG CCC CAG GDN BAA CAT GCA CTG CCT TAC CTT CGC GAT ATC CCC AGG CGT AAT TCT |
| MS_AmpC_S228_2_p6 | T*G*G* TCG ACT TCA CAC CAT AAG CTT CAG CAT CTA ACG CCC CAG GDH NAA CAT GCA CTG CCT TAC CTT CGC GAT ATC CCC AGG CGT AAT TCT |
| MS_AmpC_P229_1_p6 | C*A*A* TGG TCG ACT TCA CAC CAT AAG CTT CAG CAT CTA ACG CCC CBN HCG AAA CAT GCA CTG CCT TAC CTT CGC GAT ATC CCC AGG CGT AAT |
| MS_AmpC_P229_2_p6 | C*A*A* TGG TCG ACT TCA CAC CAT AAG CTT CAG CAT CTA ACG CCC CBH NCG AAA CAT GCA CTG CCT TAC CTT CGC GAT ATC CCC AGG CGT AAT |
| MS_AmpC_G230_1_p6 | C*T*T* CAA TGG TCG ACT TCA CAC CAT AAG CTT CAG CAT CTA ACG CDN DAG GCG AAA CAT GCA CTG CCT TAC CTT CGC GAT ATC CCC AGG CGT |
| MS_AmpC_G230_2_p6 | C*T*T* CAA TGG TCG ACT TCA CAC CAT AAG CTT CAG CAT CTA ACG CDD NAG GCG AAA CAT GCA CTG CCT TAC CTT CGC GAT ATC CCC AGG CGT |
| MS_AmpC_A231_1_p6 | T*A*T* CTT CAA TGG TCG ACT TCA CAC CAT AAG CTT CAG CAT CTA ADN DCC CAG GCG AAA CAT GCA CTG CCT TAC CTT CGC GAT ATC CCC AGG |
| MS_AmpC_A231_2_p6 | T*A*T* CTT CAA TGG TCG ACT TCA CAC CAT AAG CTT CAG CAT CTA ADH NCC CAG GCG AAA CAT GCA CTG CCT TAC CTT CGC GAT ATC CCC AGG |
| MS_AmpC_L232_1_p6 | C*C*A* TAT CTT CAA TGG TCG ACT TCA CAC CAT AAG CTT CAG CAT CVN BCG CCC CAG GCG AAA CAT GCA CTG CCT TAC CTT CGC GAT ATC CCC |
| MS_AmpC_L232_2_p6 | C*C*A* TAT CTT CAA TGG TCG ACT TCA CAC CAT AAG CTT CAG CAT CVB NCG CCC CAG GCG AAA CAT GCA CTG CCT TAC CTT CGC GAT ATC CCC |
| MS_AmpC_D233_1_p6 | G*G*G* CCA TAT CTT CAA TGG TCG ACT TCA CAC CAT AAG CTT CAG CBN DTA ACG CCC CAG GCG AAA CAT GCA CTG CCT TAC CTT CGC GAT ATC |
| MS_AmpC_D233_2_p6 | G*G*G* CCA TAT CTT CAA TGG TCG ACT TCA CAC CAT AAG CTT CAG CBV NTA ACG CCC CAG GCG AAA CAT GCA CTG CCT TAC CTT CGC GAT ATC |
| MS_AmpC_A234_1_p6 | A*G*C* GGG CCA TAT CTT CAA TGG TCG ACT TCA CAC CAT AAG CTT CBN DAT CTA ACG CCC CAG GCG AAA CAT GCA CTG CCT TAC CTT CGC GAT |
| MS_AmpC_A234_2_p6 | A*G*C* GGG CCA TAT CTT CAA TGG TCG ACT TCA CAC CAT AAG CTT CBH NAT CTA ACG CCC CAG GCG AAA CAT GCA CTG CCT TAC CTT CGC GAT |

| | |
|---|---|
| MS_AmpC_E235_1_p6 | C*C*C* AGC GGG CCA TAT CTT CAA TGG TCG ACT TCA CAC CAT AAG CVN DAG CAT CTA ACG CCC CAG GCG AAA CAT GCA CTG CCT TAC CTT CGC |
| MS_AmpC_E235_2_p6 | C*C*C* AGC GGG CCA TAT CTT CAA TGG TCG ACT TCA CAC CAT AAG CVV NAG CAT CTA ACG CCC CAG GCG AAA CAT GCA CTG CCT TAC CTT CGC |
| MS_AmpC_A236_1_p6 | G*C*A* CCC AGC GGG CCA TAT CTT CAA TGG TCG ACT TCA CAC CAT ABN DTT CAG CAT CTA ACG CCC CAG GCG AAA CAT GCA CTG CCT TAC CTT |
| MS_AmpC_A236_2_p6 | G*C*A* CCC AGC GGG CCA TAT CTT CAA TGG TCG ACT TCA CAC CAT ABH NTT CAG CAT CTA ACG CCC CAG GCG AAA CAT GCA CTG CCT TAC CTT |
| MS_AmpC_Y237_1_p6 | T*T*T* GCA CCC AGC GGG CCA TAT CTT CAA TGG TCG ACT TCA CAC CBN BAG CTT CAG CAT CTA ACG CCC CAG GCG AAA CAT GCA CTG CCT TAC |
| MS_AmpC_Y237_2_p6 | T*T*T* GCA CCC AGC GGG CCA TAT CTT CAA TGG TCG ACT TCA CAC CBV NAG CTT CAG CAT CTA ACG CCC CAG GCG AAA CAT GCA CTG CCT TAC |
| MS_AmpC_G238_1_p6 | T*G*C* TTT GCA CCC AGC GGG CCA TAT CTT CAA TGG TCG ACT TCA CBN DAT AAG CTT CAG CAT CTA ACG CCC CAG GCG AAA CAT GCA CTG CCT |
| MS_AmpC_G238_2_p6 | T*G*C* TTT GCA CCC AGC GGG CCA TAT CTT CAA TGG TCG ACT TCA CBD NAT AAG CTT CAG CAT CTA ACG CCC CAG GCG AAA CAT GCA CTG CCT |
| MS_AmpC_K240_1_p6 | T*T*A* AAT TGC TTT GCA CCC AGC GGG CCA TAT CTT CAA TGG TCG ADN VCA CAC CAT AAG CTT CAG CAT CTA ACG CCC CAG GCG AAA CAT GCA |
| MS_AmpC_K240_2_p6 | T*T*A* AAT TGC TTT GCA CCC AGC GGG CCA TAT CTT CAA TGG TCG ADV NCA CAC CAT AAG CTT CAG CAT CTA ACG CCC CAG GCG AAA CAT GCA |
| MS_AmpC_E288_1_p7 | T*G*C* CGT TAA TGA TGC TGT CAG GAT TTA CCG GCC AGT CCA GCA TVN DCC AGC CCA GGC CCT GAT ACA TAT CGC CGG TTT GCC AGT AGC GAG |
| MS_AmpC_E288_2_p7 | T*G*C* CGT TAA TGA TGC TGT CAG GAT TTA CCG GCC AGT CCA GCA TVV NCC AGC CCA GGC CCT GAT ACA TAT CGC CGG TTT GCC AGT AGC GAG |
| MS_AmpC_S303_1_p8 | T*A*A* TCG CTT TTA CGG GGC GTG CTG CCA GTG CAA TTT TAT TGT CBN VGC CGT TAA TGA TGC TGT CAG GAT TTA CCG GCC AGT CCA GCA TTT |
| MS_AmpC_S303_2_p8 | T*A*A* TCG CTT TTA CGG GGC GTG CTG CCA GTG CAA TTT TAT TGT CBD NGC CGT TAA TGA TGC TGT CAG GAT TTA CCG GCC AGT CCA GCA TTT |
| MS_AmpC_D304_1_p8 | G*C*G* TAA TCG CTT TTA CGG GGC GTG CTG CCA GTG CAA TTT TAT THN DAC TGC CGT TAA TGA TGC TGT CAG GAT TTA CCG GCC AGT CCA GCA |
| MS_AmpC_D304_2_p8 | G*C*G* TAA TCG CTT TTA CGG GGC GTG CTG CCA GTG CAA TTT TAT THV NAC TGC CGT TAA TGA TGC TGT CAG GAT TTA CCG GCC AGT CCA GCA |

| | |
|---|---|
| MS_AmpC_N305_1_p8 | G*G*G* GCG TAA TCG CTT TTA CGG GGC GTG CTG CCA GTG CAA TTT TBN VGT CAC TGC CGT TAA TGA TGC TGT CAG GAT TTA CCG GCC AGT CCA |
| MS_AmpC_N305_2_p8 | G*G*G* GCG TAA TCG CTT TTA CGG GGC GTG CTG CCA GTG CAA TTT TBV NGT CAC TGC CGT TAA TGA TGC TGT CAG GAT TTA CCG GCC AGT CCA |
| MS_AmpC_I307_1_p8 | G*A*G* TTG GGG GCG TAA TCG CTT TTA CGG GGC GTG CTG CCA GTG CBN VTT TAT TGT CAC TGC CGT TAA TGA TGC TGT CAG GAT TTA CCG GCC |
| MS_AmpC_I307_2_p8 | G*A*G* TTG GGG GCG TAA TCG CTT TTA CGG GGC GTG CTG CCA GTG CBB NTT TAT TGT CAC TGC CGT TAA TGA TGC TGT CAG GAT TTA CCG GCC |
| MS_AmpC_A308_1_p8 | C*A*G* GAG TTG GGG GCG TAA TCG CTT TTA CGG GGC GTG CTG CCA GVN DAA TTT TAT TGT CAC TGC CGT TAA TGA TGC TGT CAG GAT TTA CCG |
| MS_AmpC_A308_2_p8 | C*A*G* GAG TTG GGG GCG TAA TCG CTT TTA CGG GGC GTG CTG CCA GVH NAA TTT TAT TGT CAC TGC CGT TAA TGA TGC TGT CAG GAT TTA CCG |
| MS_AmpC_L309_1_p8 | C*T*G* CAG GAG TTG GGG GCG TAA TCG CTT TTA CGG GGC GTG CTG CDN HTG CAA TTT TAT TGT CAC TGC CGT TAA TGA TGC TGT CAG GAT TTA |
| MS_AmpC_L309_2_p8 | C*T*G* CAG GAG TTG GGG GCG TAA TCG CTT TTA CGG GGC GTG CTG CDB NTG CAA TTT TAT TGT CAC TGC CGT TAA TGA TGC TGT CAG GAT TTA |
| MS_AmpC_A310_1_p8 | G*T*A* CTG CAG GAG TTG GGG GCG TAA TCG CTT TTA CGG GGC GTG CVN DCA GTG CAA TTT TAT TGT CAC TGC CGT TAA TGA TGC TGT CAG GAT |
| MS_AmpC_A310_2_p8 | G*T*A* CTG CAG GAG TTG GGG GCG TAA TCG CTT TTA CGG GGC GTG CVH NCA GTG CAA TTT TAT TGT CAC TGC CGT TAA TGA TGC TGT CAG GAT |
| MS_AmpC_R312_1_p8 | A*T*G* CGC GTA CTG CAG GAG TTG GGG GCG TAA TCG CTT TTA CGG GHN HTG CTG CCA GTG CAA TTT TAT TGT CAC TGC CGT TAA TGA TGC TGT |
| MS_AmpC_R312_2_p8 | A*T*G* CGC GTA CTG CAG GAG TTG GGG GCG TAA TCG CTT TTA CGG GHD NTG CTG CCA GTG CAA TTT TAT TGT CAC TGC CGT TAA TGA TGC TGT |
| MS_AmpC_H330_1_p9 | T*A*A* ACG CGA CAT AGC TAC CAA ATC CGC CGG TCG CCC CTG TTT TBN HTA CCC ATG ATG CGC GTA CTG CAG GAG TTG GGG GCG TAA TCG CTT |
| MS_AmpC_H330_2_p9 | T*A*A* ACG CGA CAT AGC TAC CAA ATC CGC CGG TCG CCC CTG TTT TBV NTA CCC ATG ATG CGC GTA CTG CAG GAG TTG GGG GCG TAA TCG CTT |
| MS_AmpC_K331_1_p9 | G*A*A* TAA ACG CGA CAT AGC TAC CAA ATC CGC CGG TCG CCC CTG TVN VAT GTA CCC ATG ATG CGC GTA CTG CAG GAG TTG GGG GCG TAA TCG |
| MS_AmpC_K331_2_p9 | G*A*A* TAA ACG CGA CAT AGC TAC CAA ATC CGC CGG TCG CCC CTG TVV NAT GTA CCC ATG ATG CGC GTA CTG CAG GAG TTG GGG GCG TAA TCG |

| | |
|---|---|
| MS_AmpC_T332_1_p9 | C*T*G* GAA TAA ACG CGA CAT AGC TAC CAA ATC CGC CGG TCG CCC CVN VTT TAT GTA CCC ATG ATG CGC GTA CTG CAG GAG TTG GGG GCG TAA |
| MS_AmpC_T332_2_p9 | C*T*G* GAA TAA ACG CGA CAT AGC TAC CAA ATC CGC CGG TCG CCC CVH NTT TAT GTA CCC ATG ATG CGC GTA CTG CAG GAG TTG GGG GCG TAA |
| MS_AmpC_G333_1_p9 | T*T*T* CTG GAA TAA ACG CGA CAT AGC TAC CAA ATC CGC CGG TCG CDN DTG TTT TAT GTA CCC ATG ATG CGC GTA CTG CAG GAG TTG GGG GCG |
| MS_AmpC_G333_2_p9 | T*T*T* CTG GAA TAA ACG CGA CAT AGC TAC CAA ATC CGC CGG TCG CDD NTG TTT TAT GTA CCC ATG ATG CGC GTA CTG CAG GAG TTG GGG GCG |
| MS_AmpC_A334_1_p9 | C*T*T* TTT CTG GAA TAA ACG CGA CAT AGC TAC CAA ATC CGC CGG TDN DCC CTG TTT TAT GTA CCC ATG ATG CGC GTA CTG CAG GAG TTG GGG |
| MS_AmpC_A334_2_p9 | C*T*T* TTT CTG GAA TAA ACG CGA CAT AGC TAC CAA ATC CGC CGG TDH NCC CTG TTT TAT GTA CCC ATG ATG CGC GTA CTG CAG GAG TTG GGG |
| MS_AmpC_T335_1_p9 | G*C*T* CTT TTT CTG GAA TAA ACG CGA CAT AGC TAC CAA ATC CGC CHN VCG CCC CTG TTT TAT GTA CCC ATG ATG CGC GTA CTG CAG GAG TTG |
| MS_AmpC_T335_2_p9 | G*C*T* CTT TTT CTG GAA TAA ACG CGA CAT AGC TAC CAA ATC CGC CHH NCG CCC CTG TTT TAT GTA CCC ATG ATG CGC GTA CTG CAG GAG TTG |
| MS_AmpC_N362_1_p10 | G*T*A* GAG CGT TAA GAA TCT GCC AGG CGG CGT CGA CTC TCG CTG GBN VGG GAT AGT TTT TGT TTG CCA GCA TCA CGA TAC CCA GCT CTT TTT |
| MS_AmpC_N362_2_p10 | G*T*A* GAG CGT TAA GAA TCT GCC AGG CGG CGT CGA CTC TCG CTG GBV NGG GAT AGT TTT TGT TTG CCA GCA TCA CGA TAC CCA GCT CTT TTT |
| MS_AmpC_R365_1_p10 | A*A*T* TTT ACT GTA GAG CGT TAA GAA TCT GCC AGG CGG CGT CGA CVN VCG CTG GAT TGG GAT AGT TTT TGT TTG CCA GCA TCA CGA TAC CCA |
| MS_AmpC_R365_2_p10 | A*A*T* TTT ACT GTA GAG CGT TAA GAA TCT GCC AGG CGG CGT CGA CVD NCG CTG GAT TGG GAT AGT TTT TGT TTG CCA GCA TCA CGA TAC CCA |

**Supplementary Table 2. Intragenic mutations in the *ampC* gene alter bacterial resistance to avibactam**

| Mutant | $IC50_{PIP}$ ($\mu$g/ml) | $IC50_{PIP}\|^{AVI}$ ($\mu$g/ml) | $IC50_{ATM}$ ($\mu$g/ml) | $IC50_{ATM}\|^{AVI}$ ($\mu$g/ml) | $IC50_{FEP}$ ($\mu$g/ml) | $IC50_{FEP}\|^{AVI}$ ($\mu$g/ml) |
|---|---|---|---|---|---|---|
| R148D | 1.9E+00 | 7.9E-01 | 1.8E+00 | 1.2E+00 | 4.1E-01 | 3.6E-02 |
| R148E | 2.7E+00 | 8.3E-01 | 4.7E+00 | 9.8E-01 | 1.3E-01 | 2.0E-02 |
| R148T | 1.7E+00 | 4.8E-01 | 3.4E+00 | 9.6E-01 | 2.6E-01 | 4.0E-02 |
| R148N | 1.7E+00 | 6.6E-01 | 1.7E+00 | 1.0E+00 | 1.7E-01 | 1.4E-02 |
| R148F | 2.9E+00 | 1.0E+00 | 3.9E+00 | 1.3E+00 | 1.2E-01 | 2.8E-02 |
| R148W | 5.0E+00 | 4.3E-01 | 1.9E+00 | 7.9E-01 | 8.5E-01 | 5.3E-02 |
| R148P | 3.9E+00 | 9.6E-01 | 1.3E+00 | 6.3E-01 | 1.9E+00 | 3.2E-01 |
| Y150R | 8.3E+00 | 6.3E+00 | 1.9E+01 | 1.3E+01 | 1.4E-02 | 1.4E-02 |
| Y150K | 1.0E+01 | 1.2E+01 | 2.1E+01 | 2.0E+01 | 1.4E-02 | 1.4E-02 |
| Y150S | 8.3E+00 | 2.7E+00 | 2.4E+01 | 1.9E+01 | 1.4E-02 | 1.9E-02 |
| Y150T | 9.2E+00 | 4.3E+00 | 9.2E+00 | 8.4E+00 | 1.4E-02 | 1.4E-02 |
| Y150N | 3.0E+00 | 2.4E+00 | 8.5E+00 | 4.5E+00 | 2.8E-02 | 1.4E-02 |
| Y150Q | 4.7E+00 | 3.3E+00 | 4.2E+00 | 4.0E+00 | 1.4E-02 | 1.4E-02 |
| Y150A | 9.1E+00 | 3.4E+00 | 5.0E+01 | 4.0E+01 | 1.4E-02 | 1.4E-02 |
| Y150V | 7.8E+00 | 3.2E+00 | 3.8E+00 | 3.4E+00 | 1.4E-02 | 1.4E-02 |
| Y150I | 5.1E+00 | 6.1E+00 | 8.6E+00 | 3.9E+00 | 1.4E-02 | 1.4E-02 |
| Y150L | 7.1E+00 | 6.0E+00 | 4.6E+00 | 3.6E+00 | 1.4E-02 | 1.4E-02 |
| Y150M | 8.8E+00 | 7.3E+00 | 4.6E+00 | 3.6E+00 | 1.4E-02 | 1.4E-02 |
| Y150C | 7.3E+00 | 3.0E+00 | 9.4E+00 | 8.3E+00 | 1.4E-02 | 1.4E-02 |
| Y150G | 1.1E+01 | 6.5E+00 | 1.8E+01 | 9.5E+00 | 1.4E-02 | 1.4E-02 |
| Y150P | 3.8E+00 | 2.3E+00 | 2.4E+01 | 2.0E+01 | 1.4E-02 | 1.4E-02 |
| N152S | 5.3E+01 | 2.6E+00 | 1.2E+00 | 1.1E+00 | 1.4E-02 | 1.4E-02 |
| N152T | 4.9E+01 | 6.6E+00 | 1.4E+00 | 7.5E-01 | 1.4E-02 | 1.4E-02 |
| N152Q | 7.2E+00 | 2.4E+00 | 6.3E-01 | 1.1E+00 | 1.4E-02 | 1.4E-02 |
| N152A | 3.1E+00 | 2.6E+00 | 1.5E+00 | 3.6E+00 | 1.4E-02 | 1.4E-02 |
| N152G | 2.2E+01 | 2.3E+00 | 6.3E-01 | 7.6E-01 | 1.4E-02 | 1.4E-02 |
| E272R | 2.4E+01 | 7.5E-01 | 1.8E+01 | 6.3E-01 | 4.6E-02 | 2.6E-02 |
| E272K | 1.3E+01 | 8.5E-01 | 1.5E+01 | 1.2E+00 | 3.8E-02 | 2.5E-02 |
| E272S | 2.2E+01 | 3.9E-01 | 7.3E+00 | 6.3E-01 | 2.9E-02 | 2.3E-02 |
| E272T | 2.0E+01 | 6.4E-01 | 8.2E+00 | 6.3E-01 | 3.1E-02 | 2.5E-02 |
| E272N | 1.2E+01 | 2.9E-01 | 7.8E+00 | 1.4E+00 | 3.2E-02 | 4.4E-02 |
| E272I | 2.2E+01 | 6.8E-01 | 1.4E+01 | 6.3E-01 | 6.5E-02 | 2.7E-02 |
| E272L | 1.1E+01 | 6.8E-01 | 9.9E+00 | 6.3E-01 | 1.4E-02 | 1.4E-02 |
| E272F | 2.4E+01 | 8.7E-01 | 4.6E+00 | 6.3E-01 | 3.0E-02 | 2.0E-02 |
| E272Y | 1.9E+01 | 8.1E-01 | 4.6E+00 | 6.3E-01 | 4.1E-02 | 1.5E-02 |
| E272C | 2.3E+01 | 6.6E-01 | 4.3E+00 | 1.3E+00 | 3.6E-02 | 2.4E-02 |
| S287R | 4.8E+01 | 6.7E-01 | 3.7E+00 | 6.3E-01 | 5.2E-01 | 1.4E-02 |
| S287H | 2.0E+01 | 7.0E-01 | 5.1E+00 | 6.3E-01 | 3.8E+00 | 8.8E-02 |
| S287K | 4.1E+01 | 5.5E-01 | 3.4E+00 | 6.3E-01 | 4.5E-01 | 1.4E-02 |
| S287D | 1.1E+01 | 8.1E-01 | 1.9E+00 | 6.3E-01 | 2.2E+00 | 7.0E-02 |

| | | | | | | |
|---|---|---|---|---|---|---|
| S287E | 1.7E+00 | 7.0E-01 | 2.3E+00 | 6.3E-01 | 3.7E-01 | 4.6E-02 |
| S287T | 1.4E+01 | 6.6E-01 | 4.1E+00 | 1.2E+00 | 4.2E-01 | 3.7E-02 |
| S287N | 1.2E+01 | 6.0E-01 | 4.4E+00 | 1.1E+00 | 8.9E-01 | 2.4E-02 |
| S287V | 2.0E+01 | 6.9E-01 | 6.9E+00 | 6.3E-01 | 2.0E-01 | 2.8E-02 |
| S287I | 1.7E+01 | 7.4E-01 | 3.9E+00 | 6.3E-01 | 6.1E-01 | 3.9E-02 |
| S287L | 1.1E+01 | 8.4E-01 | 4.5E+00 | 6.3E-01 | 1.6E+00 | 4.3E-02 |
| S287M | 1.0E+01 | 6.1E-01 | 4.1E+00 | 6.3E-01 | 4.8E-01 | 2.7E-02 |
| S287F | 1.8E+01 | 6.7E-01 | 2.9E+00 | 6.3E-01 | 2.2E-01 | 1.4E-02 |
| S287Y | 2.1E+01 | 6.7E-01 | 1.8E+00 | 6.3E-01 | 2.6E-01 | 2.5E-02 |
| S287W | 2.6E+01 | 6.5E-01 | 2.9E+00 | 6.3E-01 | 5.7E-01 | 1.4E-02 |
| S287G | 4.2E+00 | 6.5E-01 | 4.6E+00 | 6.3E-01 | 9.3E-01 | 6.7E-02 |
| S287P | 2.5E+01 | 7.4E-01 | 3.7E+00 | 6.3E-01 | 6.2E+00 | 3.5E-02 |
| D288R | 1.3E+01 | 6.8E-01 | 4.8E+00 | 6.3E-01 | 2.8E-02 | 2.5E-02 |
| D288T | 2.1E+01 | 2.3E-01 | 3.6E+00 | 6.3E-01 | 1.4E-02 | 1.4E-02 |
| D288Q | 2.3E+01 | 6.6E-01 | 5.9E+00 | 1.3E+00 | 3.9E-02 | 2.5E-02 |
| D288G | 2.3E+01 | 7.2E-01 | 4.3E+00 | 7.2E-01 | 3.6E-02 | 1.8E-02 |
| D288P | 4.1E+00 | 7.2E-01 | 3.1E+00 | 6.9E-01 | 5.1E-01 | 4.6E-02 |
| N289R | 1.6E+01 | 4.1E-01 | 3.2E+00 | 6.3E-01 | 1.9E-02 | 1.4E-02 |
| N289S | 2.5E+01 | 2.6E-01 | 4.5E+00 | 7.2E-01 | 2.4E-02 | 1.4E-02 |
| N289T | 3.0E+01 | 5.6E-01 | 3.5E+00 | 1.3E+00 | 1.4E-02 | 1.4E-02 |
| N289Q | 2.6E+01 | 2.0E-01 | 4.0E+00 | 8.3E-01 | 1.4E-02 | 1.4E-02 |
| N289A | 3.0E+01 | 1.9E-01 | 4.0E+00 | 6.3E-01 | 2.4E-02 | 1.4E-02 |
| N289V | 4.6E+01 | 6.2E-01 | 1.8E+00 | 6.3E-01 | 1.4E-02 | 1.4E-02 |
| N289I | 7.6E+01 | 3.2E-01 | 1.6E+00 | 6.3E-01 | 1.4E-02 | 1.4E-02 |
| N289L | 2.9E+01 | 1.9E-01 | 4.1E+00 | 1.2E+00 | 1.4E-02 | 1.4E-02 |
| N289M | 5.7E+01 | 7.1E-01 | 3.9E+00 | 6.3E-01 | 3.9E-02 | 1.4E-02 |
| N289F | 2.2E+01 | 6.2E-01 | 1.6E+00 | 6.3E-01 | 1.4E-02 | 1.4E-02 |
| N289W | 2.5E+01 | 2.1E-01 | 1.5E+00 | 6.3E-01 | 1.4E-02 | 1.4E-02 |
| N289G | 2.5E+01 | 6.6E-01 | 8.1E+00 | 6.3E-01 | 1.1E-01 | 3.9E-02 |
| N289P | 3.7E+01 | 1.9E-01 | 3.7E+00 | 6.3E-01 | 2.5E-02 | 1.4E-02 |
| I291D | 4.8E+00 | 8.1E-01 | 4.1E+00 | 6.8E-01 | 1.2E+00 | 6.3E-02 |
| I291A | 1.1E+01 | 5.6E-01 | 3.6E+00 | 6.3E-01 | 5.6E-02 | 2.8E-02 |
| I291L | 6.0E+00 | 1.9E-01 | 1.5E+00 | 1.1E+00 | 1.4E-02 | 1.4E-02 |
| I291G | 2.1E+01 | 7.9E-01 | 2.8E+00 | 1.3E+00 | 3.8E-01 | 1.7E-02 |
| I291P | 2.5E+01 | 6.0E-01 | 4.1E+00 | 9.1E-01 | 9.6E-01 | 2.7E-02 |
| A292R | 1.7E+00 | 5.3E-01 | 8.7E+00 | 3.3E+00 | 2.2E-02 | 1.9E-02 |
| A292K | 2.8E+01 | 7.4E-01 | 4.1E+00 | 1.4E+00 | 4.4E-02 | 1.9E-02 |
| A292T | 2.7E+01 | 5.8E-01 | 5.2E+00 | 6.3E-01 | 7.6E-02 | 1.4E-02 |
| A292Q | 2.5E+01 | 7.9E-01 | 4.0E+00 | 1.3E+00 | 1.4E-02 | 2.6E-02 |
| A292V | 3.2E+01 | 7.0E-01 | 4.2E+00 | 6.3E-01 | 6.9E-02 | 1.9E-02 |
| A292I | 1.5E+01 | 7.0E-01 | 3.0E+00 | 6.3E-01 | 3.3E-02 | 1.4E-02 |
| A292L | 1.5E+01 | 7.1E-01 | 4.0E+00 | 1.2E+00 | 3.6E-02 | 1.4E-02 |
| A292M | 1.8E+01 | 7.8E-01 | 3.5E+00 | 6.3E-01 | 1.4E-02 | 1.4E-02 |
| A292G | 3.9E+00 | 1.9E-01 | 6.1E+00 | 1.2E+00 | 8.7E-01 | 6.2E-02 |
| A292P | 3.3E+01 | 1.1E+00 | 3.8E+00 | 6.3E-01 | 8.1E-01 | 3.3E-02 |
| M293S | 1.7E+00 | 1.0E+00 | 4.0E+00 | 1.5E+00 | 2.2E-01 | 3.5E-02 |
| M293T | 1.3E+01 | 9.3E-01 | 2.7E+00 | 6.7E-01 | 3.3E-02 | 3.1E-02 |
| M293I | 1.9E+01 | 6.6E-01 | 3.5E+00 | 7.7E-01 | 1.1E-01 | 3.6E-02 |

| | | | | | | |
|---|---|---|---|---|---|---|
| M293G | 3.7E+00 | 6.7E-01 | 2.3E+00 | 1.3E+00 | 3.8E-01 | 2.7E-02 |
| M293P | 1.2E+01 | 6.4E-01 | 3.9E+00 | 9.7E-01 | 1.0E+00 | 3.1E-02 |
| A294Y | 5.1E+00 | 5.2E-01 | 6.1E+00 | 6.3E-01 | 5.9E-01 | 5.2E-02 |
| R296E | 3.8E+00 | 5.9E-01 | 2.8E+00 | 6.3E-01 | 2.9E-01 | 3.7E-02 |
| R296P | 1.2E+01 | 7.6E-01 | 3.6E+00 | 6.3E-01 | 3.5E+00 | 4.3E-02 |
| H314I | 1.0E+01 | 1.1E+00 | 3.8E+00 | 9.7E-01 | 7.4E-02 | 1.7E-02 |
| H314L | 4.8E+00 | 1.0E+00 | 4.9E+00 | 1.4E+00 | 1.1E-01 | 4.9E-02 |
| K315Y | 1.7E+00 | 1.1E+00 | 1.5E+00 | 1.4E+00 | 2.3E-02 | 3.1E-02 |
| T316R | 2.2E+01 | 8.6E-01 | 7.2E+00 | 3.2E+00 | 1.4E-02 | 2.2E-02 |
| T316V | 1.7E+00 | 7.3E-01 | 1.8E+00 | 1.3E+00 | 1.4E-02 | 1.4E-02 |
| T316W | 1.7E+00 | 8.2E-01 | 1.7E+00 | 3.2E+00 | 1.4E-02 | 1.4E-02 |
| T316G | 4.8E+00 | 7.2E-01 | 3.7E+00 | 1.3E+00 | 2.5E-02 | 2.4E-02 |
| A318G | 1.9E+01 | 1.0E+00 | 3.9E+00 | 7.4E-01 | 1.4E-02 | 2.2E-02 |
| T319I | 1.7E+00 | 7.2E-01 | 3.9E+00 | 1.9E+00 | 1.4E-02 | 1.4E-02 |
| N346R | 3.8E+00 | 8.0E-01 | 5.1E+00 | 1.6E+00 | 8.8E-02 | 3.4E-02 |
| N346H | 1.7E+00 | 1.0E+00 | 3.5E+00 | 1.9E+00 | 3.3E-02 | 1.4E-02 |
| N346E | 1.7E+00 | 7.9E-01 | 1.2E+00 | 6.3E-01 | 2.9E-02 | 2.2E-02 |
| N346S | 6.5E+00 | 1.0E+00 | 4.0E+00 | 1.4E+00 | 3.0E-02 | 2.0E-02 |
| N346T | 1.0E+01 | 1.0E+00 | 3.4E+00 | 1.1E+00 | 3.0E-02 | 1.4E-02 |
| N346A | 6.7E+00 | 1.3E+00 | 4.0E+00 | 3.0E+00 | 2.3E-02 | 1.6E-02 |
| N346V | 8.9E+00 | 1.1E+00 | 5.4E+00 | 1.6E+00 | 1.1E-01 | 3.7E-02 |
| N346I | 4.9E+00 | 9.6E-01 | 2.3E+00 | 6.3E-01 | 5.6E-02 | 2.8E-02 |
| N346L | 4.5E+00 | 1.0E+00 | 3.5E+00 | 6.3E-01 | 2.4E-01 | 4.5E-02 |
| N346M | 1.7E+00 | 1.1E+00 | 3.5E+00 | 1.4E+00 | 3.4E-02 | 1.4E-02 |
| N346F | 1.7E+00 | 1.4E+00 | 3.1E+00 | 1.6E+00 | 1.2E-01 | 9.2E-02 |
| N346Y | 1.7E+00 | 1.4E+00 | 3.9E+00 | 3.8E+00 | 6.3E-01 | 1.1E+00 |
| N346W | 4.9E+00 | 1.5E+00 | 3.5E+00 | 1.7E+00 | 1.9E+00 | 5.7E-01 |
| N346C | 1.7E+00 | 8.5E-01 | 2.9E+00 | 1.3E+00 | 2.7E-02 | 2.5E-02 |
| N346G | 8.4E+00 | 1.2E+00 | 4.2E+00 | 3.7E+00 | 2.9E-02 | 2.4E-02 |
| N346P | 4.1E+00 | 1.1E+00 | 3.4E+00 | 1.4E+00 | 1.4E-02 | 1.8E-02 |
| R349L | 4.6E+00 | 1.1E+00 | 1.3E+00 | 7.4E-01 | 3.2E-02 | 4.2E-02 |
| R349F | 5.1E+00 | 1.1E+00 | 2.1E+00 | 8.1E-01 | 4.7E-02 | 2.6E-02 |
| R349Y | 1.7E+00 | 1.0E+00 | 1.6E+00 | 6.3E-01 | 9.9E-02 | 3.7E-02 |
| R349P | 4.6E+00 | 1.0E+00 | 2.3E+00 | 6.3E-01 | 8.3E-02 | 3.0E-02 |
| WT | 6.7E+00 | 1.0E+00 | 8.7E+00 | 1.1E+00 | 4.7E-02 | 3.4E-02 |
